# New Additions to the CRISPR Toolbox: CRISPR-*CLONInG* and CRISPR-*CLIP* for Donor Construction in Genome Editing

**DOI:** 10.1101/746826

**Authors:** Dorjee T.N. Shola, Chingwen Yang, Vhy-Shelta Kewaldar, Pradip Kar, Victor Bustos

## Abstract

CRISPR-Cas has proven to be the most versatile genetic tinkering system of our time, predominantly as a precision genome editing tool. Here, we demonstrate two additions to the repertoire of CRISPR’s application for constructing donor DNA templates. (i) CRISPR-*CLONInG* (CRISPR-*<u>C</u>*utting & *<u>L</u>*igation *<u>O</u>*f *<u>N</u>*ucleic acid *<u>In</u> vitro* via *<u>G</u>*ibson) was devised to enable efficient cut-and-paste of multiple complex DNA fragments by using CRISPR/Cas9 as a digestion alternative with precision and exclusivity features, followed by joining the digested products via Gibson Assembly, to rapidly construct dsDNA and AAV donor vectors without cloning scars. (ii) CRISPR-*CLIP* (CRISPR-*<u>C</u>*lipped *<u>L</u>*ong ssDNA via *I*ncising *<u>P</u>*lasmid) was devised as a DNA clipping tool to efficiently retrieve long single-stranded DNA (lssDNA) from plasmid, up to 3.5 kbases, which can be supplied as the donor template for creating genetically engineered mice via *Easi*-CRISPR. We utilized two different Cas types (Cpf1 and Cas9n) to induce two distinct incisions at the respective ends of the lssDNA cassette junctions on the plasmid, yielding three independent single-stranded DNA units of unique sizes eligible for strand separation, followed by target strand clip-out through gel extraction. The retrieval of the lssDNA donor circumvents involvements of restriction enzymes and DNA polymerase-based steps, hence not only retains sequence fidelity but carries virtually no restriction on sequence composition, further mitigating limitations on current *Easi*-CRISPR method. With the add-on feature of universal DNA-tag sequences of Cpf1-Cas9 duo PAM, CRISPR-*CLIP* can be facile and applicable to generate lssDNA templates for any genomic target of choice. Additionally, we demonstrate robust gene editing efficiencies in neuroblastoma cell line as well as in mice attained with the AAV and lssDNA donors constructed herein.

## Introduction

The versatility of class 2 CRISPR system is attributable to the simplicity of Cas nuclease being guided by a single programmable RNA (1), coupled with a unique spacer sequence for precise target navigation. The commonly used CRISPR protein, SpCas9, recognizes a protospacer adjacent motif (PAM) NGG, which exists once in every 42 bases in the human genome (2). In addition, the mutant version from xCas9 group recognizes an even shorter PAM ‘NG’ (3), along with Cpf1 for AT-rich sequences (4), the PAM stringency is further relaxed to allow its binding to all four nucleotides of the DNA sequence. Together with Cas9’s stability and ATP independent catalytic reaction for facilitating DNA cleavage (5), CRISPR system has attracted a wide variety of applications, prevalently as a robust tool for precision genome engineering tool in mammalian cells (6, 7).

CRISPR-mediated genome editing is carried out by using RNA guided Cas9 to induce a DNA break at the genomic locus of interest, followed by harnessing innate DNA repair mechanism to create indels for variants of gene disruptions or genetic modifications with defined outcomes. The latter is attained by additionally supplying an exogenous DNA template that carries the desired sequence flanked by homology arms (HA) to CRISPR cut site(s), and through homology-directed repair (HDR) pathway, desired modifications can be precisely integrated into the genome of target organisms in a highly efficient manner. For generating genetically engineered mouse models, donor templates can be supplied as either single-stranded oligodeoxynucleotides (ssODN) or double-stranded DNA (dsDNA) vectors (8). The former functions as an efficient donor in zygotes, yet imposes length limitations of ∼200 bases due to technical difficulties in chemical synthesis, making it suitable only for minor genetic alterations (<50 bp); whereas dsDNA-mediated editing can accommodate larger-scale genetic modifications (up to10 kb or even longer), which has been routinely conducted out via mouse embryonic stem cells (mESCs) due to poor efficiency in zygotes. Recent development of *Easi*-CRISPR (*E*fficient *a*dditions with *s*sDNA *i*nserts-CRISPR) has successfully expanded the use of single-stranded DNA donor up to ∼2 kbases long, referred to as long single-stranded DNA (lssDNA), to introduce modifications over much larger genomic regions in mouse and rat zygotes, as well as in human T cells (9–11). The length capacity of lssDNA suffices to cater to most of genome editing purposes that used to be mediated via mESC route, whereby the timeline for generating the founder mice (F0) can be accelerated to as little as two months.

The construction of dsDNA donors commonly relies on BAC recombineering or multi-step cloning methods to ligate a vector backbone and multiple DNA fragments, which oftentimes need to be acquired from various existing plasmids through the use of PCR amplification and restriction enzyme (RE). The development of seamless cloning methods, such as Gibson Assembly (12), allows multiple DNA components to be assembled into a custom donor vector in a single step *in vitro* averting any footprint and incorrect insert orientation, offering a rapid alternative to lengthy and laborious conventional approach for donor template assembly. Nonetheless, seamless cloning requires each of the assembly component to carry complementary overhang sites (e.g., type IIS RE sites and Gibson overhangs for Golden Gate and Gibson Assembly, respectively), which are routinely created by PCR amplification. Such process tends to stumble over DNA sequences with extended length or complexity, a common scenario when amplifying vector backbones with highly repetitive or palindromic sequences (e.g., multiple Lox sites in FLEx vector or ITR sequences in AAV vector), hence hindering the vector cloning. To address these PCR issues especially in acquiring vector backbones, we utilize CRISPR/Cas9 to replace RE as the digestion tool to excise undesired DNA segments from source vectors. Next, we designate Gibson Assembly over Golden Gate to facilitate seamless cloning because the former has the flexibility of only requiring overhang sites to present in either one of the DNA components to be assembled, instead of both. We term this strategy CRISPR-*CLONInG* (CRISPR-*C*utting & *L*igation of *N*ucleic acid *In vitro* via *G*ibson). Despite RE long being a vital tool for digestion, there is a very slim chance of finding a suitable candidate that cuts at the target location not only precisely, but more importantly, exclusively. For generating lssDNA donors, the current methods reckon on either PCR- or RE-based approach to procure the single-stranded templates from assembled dsDNA donors, hence the aforementioned technical challenges are categorically applicable. We wherefore demonstrate another CRISPR-based strategy, termed CRISPR-*CLIP* (CRISPR-*C*lipped *L*ong ssDNA via *I*ncising *P*lasmid) to bypass involvements of PCR and RE for lssDNA donor generation.

## Results

### CRISPR-*CLONInG* (CRISPR-*<u>C</u>*utting & *<u>L</u>*igation *<u>O</u>*f *<u>N</u>*ucleic acid *<u>In</u> vitro* via *<u>G</u>*ibson)

#### Replacing undesired DNA segment on FLEx vector with new DNA inserts

To modify a FLEx (flip-excision) donor vector, we devised CRISPR-*CLONInG* to make customizations on the existing one. We proceeded by first using two gRNAs (Luc-A & Luc-B; Table S1) with Cas9 (ctRNP complex) targeting the sites flanking the undesired Luciferase gene on the source FLEx vector for excision, rendering a backbone of 7.5 kb in length (Fig. 1A, 1D left) with complex sequences that was otherwise infeasible to acquire via inverse PCR amplification (data not shown). Next, we used PCR primers that carry corresponding 20-30 bp complementary Gibson overhangs to amplify the DNA sequences from two different plasmids for new vector inserts, FRT-Neo-FRT (∼1.87 kb) and tdTomato (∼1.43 kb) (Fig. 1B, 1D right). The three DNA components thus acquired were joined together via Gibson Assembly (Fig. 1B, 1C), with 70% success rate (Fig. 1E, SI Sequence 1). The assembled FLEx vector was transfected into mESCs and successfully achieved target modifications (data not shown). The use of CRISPR/Cas9 allows the exclusive excision of Luciferase gene from the existing vector to precisely take place at its junction sites flanked by 3’-UTR and IRES sequences (Fig. 1A), with only 1 nt deviation, which can be easily remedied by 1 extra nt carried in the primer overhangs (SI Sequence 2).

**Figure 1.**
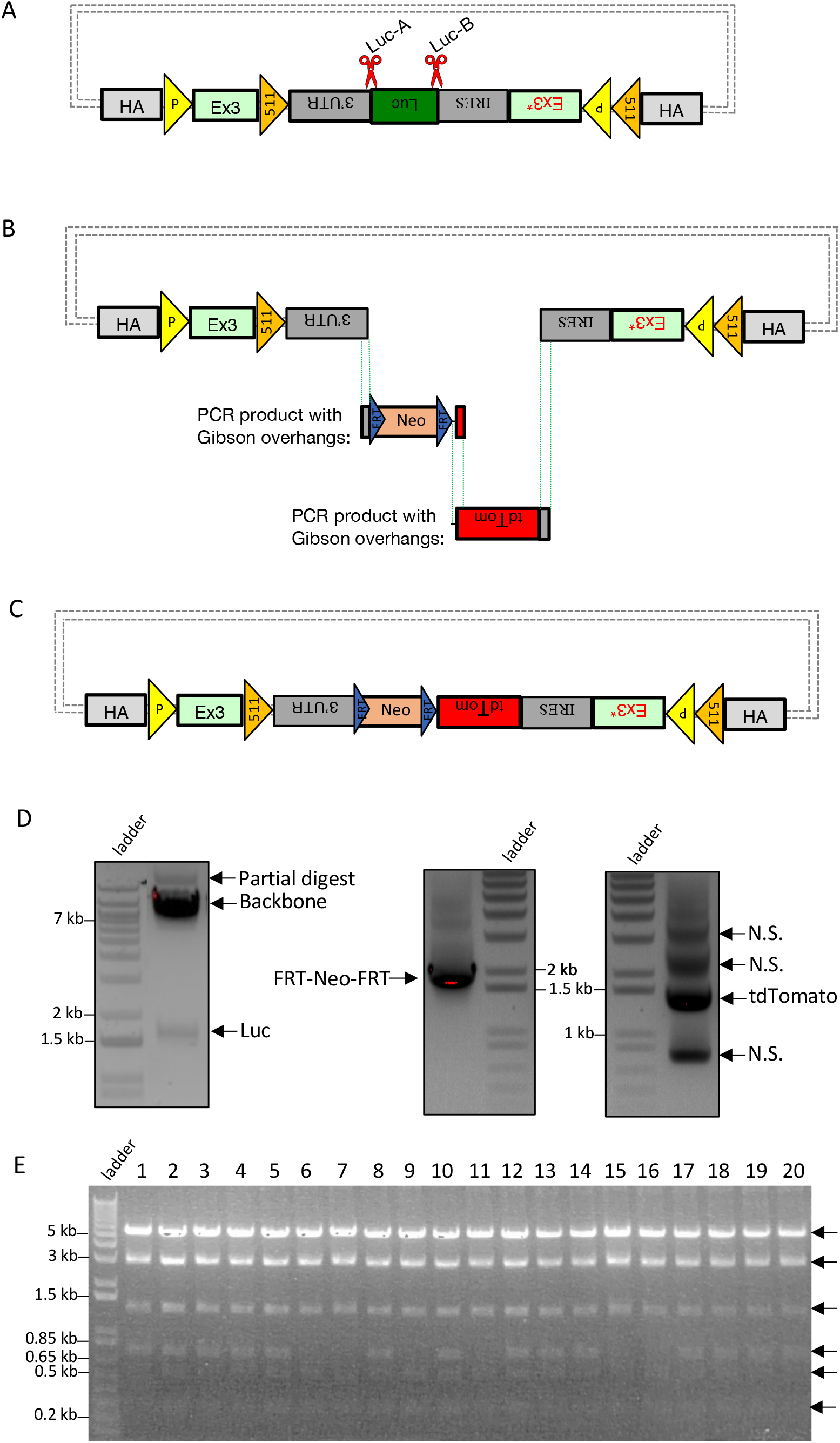
CRISPR-*CLONInG*: Replacement of Luciferase (Luc) on FLEx vector. **(A)** Schematic illustration of FLEx vector with CRISPR cut sites (red scissors) at the two junction sites flanking the undesired Luc fragment. Gray dot dashes: default backbone containing origin of replication and selection for propagation in bacterial host. **(B)** Luc was cut out with ctRNP (Cas9-ctRNA) complex; FRT-Neo-FRT and tdTomato were PCR-amplified from existing plasmids using primers carrying complementary Gibson overhangs from the adjacent DNA fragment and vector backbone. **(C)** Two new vector inserts were joined with the CRISPR-digested backbone via Gibson (HiFi) Cloning for final donor assembly. **(D)** Excised FLEx vector backbone (∼7.5 kb) and Luciferase (∼1.65 kb) (left); PCR amplified FRT-Neo-FRT (∼1.87 kb) and tdTomato (∼1.43 kb) (right). N.S.: non-specific bands. **(E)** After CRISPR-*CLONInG*, 14 out of 20 clones verified with PstI RE(s) diagnosis showed correct vector assembly (6 DNA fragments; black arrow); 3 clones validated for sequence integrity. Resolved on 0.9% agarose gel.

#### Replacing cargo sequences on AAV vector with duo-guides & gene replacement donor

Viral delivery systems serve as potent gene-delivery vehicles and have been the cornerstone of gene therapy. Adeno-associated virus (AAV) system gains preference over other viruses due to its nominal level of adverse immunogenicity in humans (13). In addition, AAV has been harnessed for targeted gene modifications (in contrast to Adeno- and Lenti-virus systems for transgenic purpose) (14) long before the advent of programmable nucleases, primarily in somatic cell lines due to its capability to effectively transduce its cargo DNA into cells which typically bear poor recombination efficiency for facilitating gene targeting (15, 16). Incorporating AAV delivery with CRISPR system has synergized the capacity of genome engineering to effectively manipulate genetic contexts in adult mice, such that disease models could be rapidly generated within several weeks and ready for biomedical studies (17, 18). While viral vector backbones are equipped with multiple cloning sites (MCS) for cargo sequence engineering, suitable MCS scarcely exists for cargos with extended length which is often the case in targeted gene editing, hence an alternative to RE-based approach is crucially needed for facilitating efficient cut-and-paste of DNA segments on an AAV vector.

Our target gene modification goal is to introduce 3 out of 5 amino acid (AA) changes within a 15-bp range (SI Sequence 3) in the exon 10 region of PSEN1 gene in neuroblastoma cell line (N2A). To construct an AAV vector to suit this purpose, we customized an existing one in two steps to replace two pieces of partial cargo DNA with desired sequences encoding CRISPR duo-guides and a donor template, respectively; the latter carries 15 bp AA replacement sequence flanked by HA to the target genomic site. The source AAV vector consists of an U6-driven guide with sgRNA cloning site, plus Cre and other components (noted as Cre-Comp). At the first stage of vector customization, we devised CRISPR-*CLONInG* using two sgRNAs (AAV-A & AAV-B; Fig. 2A, Table S1) with Cas9 (ctRNP complex) to excise Cre-Comp from AAV backbone (Fig. 2A, 2D left), followed by using PCR primers to amplify a custom gene synthesized plasmid that carries ∼0.8 kb donor template flanked with 30-35 bp Gibson overhangs (Fig. 2D right). The donor sequence was ligated into customized AAV backbone through Gibson Assembly (Fig. 2B, 2C) with >90% cloning efficiency (Fig. 2E).

**Figure 2.**
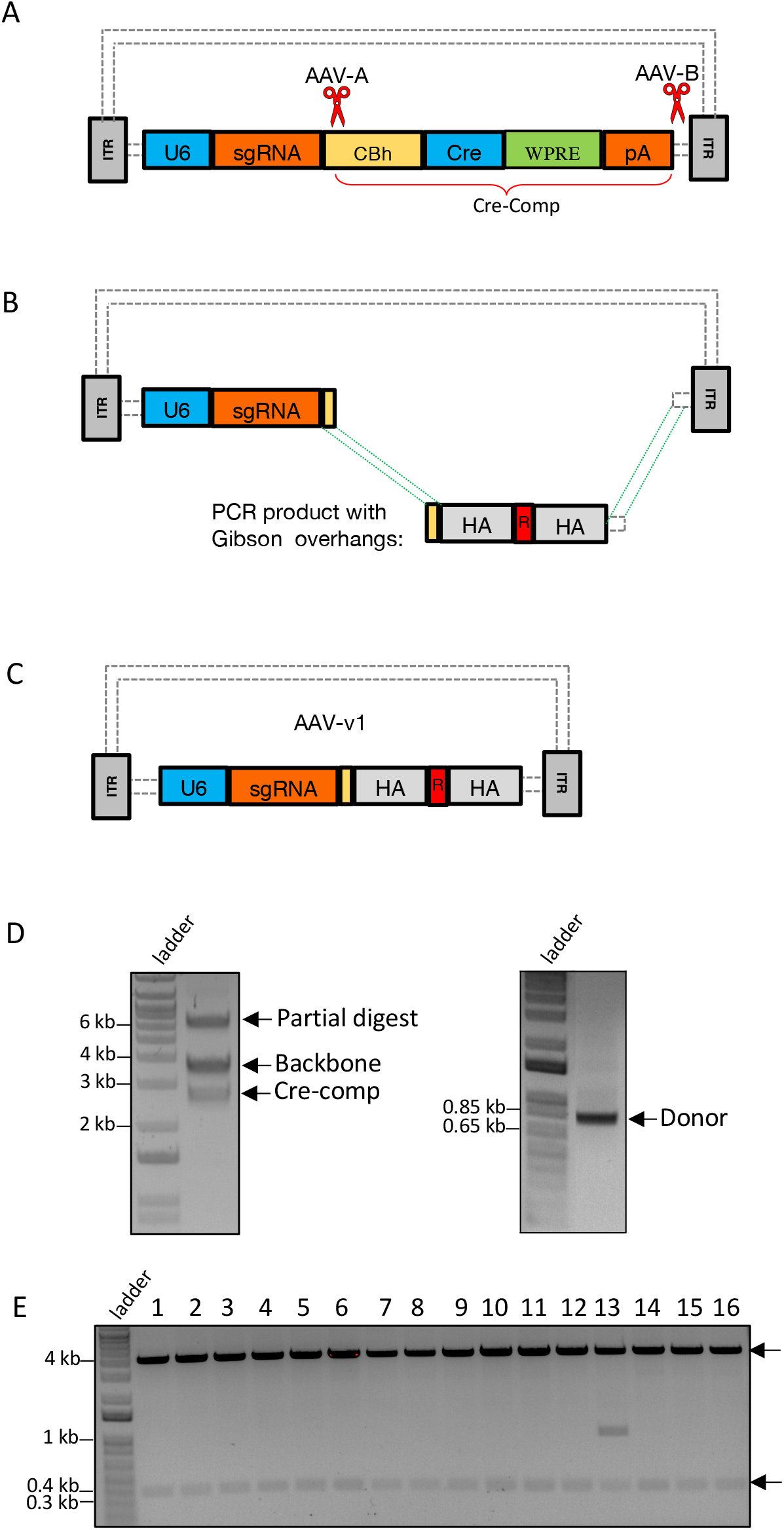
CRISPR-*CLONInG*: Replacement of partial cargo sequence (Cre-comp) with the desired donor sequence on AAV vector (Addgene #60229). **(A)** Schematic illustration of AAV vector with CRISPR cut sites (red scissors) at two ends of the Cre-comp segment. Guides (AAV-A and AAV-B) with high on-target scores were selected. **(B)** Cre-comp was cut out with ctRNP (Cas9-ctRNA) complex; donor for gene replacement (containing 15 bp AA replacement sequence, noted as ‘R’, sandwiched by HA) flanked with complementary Gibson overhangs of the adjacent AAV backbone was PCR-amplified from custom gene synthesized plasmid. **(C)** Assembled AAV-v1: donor template cloned into the customized AAV backbone via Gibson (HiFi) Assembly. **(D)** Excised AAV vector backbone (∼3.63 kb) and Cre-comp (∼2.73 kb) (left); PCR amplified donor template (∼0.8 kb) with Gibson overhangs (right). **(E)** After CRISPR-*CLONInG*, 15 out of 16 clones showed correct vector assembly, confirmed by BbsI RE(s) diagnosis (two DNA fragments; black arrow); 3 clones further validated by Sanger sequencing. Resolved on 0.9% agarose gel.

We further customized the assembled AAV vector (referred to as AAV-v1) to accommodate two CRISPR guides (sgRNA-X and sgRNA-W) targeting the exon 10 sites that span the AA change region. To exploit AAV-v1’s built-in cloning site for a single sgRNA, which upon two type-IIS enzyme (SapI) digestions, two unique 5’-overhangs (GGT on the bottom strand and GTT on the top strand) were created (Fig. 3A, middle), we designed a 0.5 kb gene synthesized plasmid carrying duo-sgRNA cassette (spacer-X+tracrRNA+U6+spacer-W) with matching overhangs, plus two uniquely positioned SapI recognition sites flanking at both ends. Specifically, one SapI recognition site was placed on the top strand 1 nt further adjacent to the overhang complementary to 5’-GGT, whereas another SapI recognition site was placed on the bottom strand 1 nt further adjacent to the overhang complementary to 5’-GTT (Fig. 3A, bottom). Upon digestions by two SapI, the aforesaid plasmid rendered two overhangs complementary to the original sgRNA cloning site on AAV-v1, which then was ligated into AAV-v1 via type-IIS RE-based cloning whereby the duo-guides were engineered into the final AAV construct (referred to as AAV-v2) (Fig. 3B). The AAV-v2 was used for viral packaging to generate the recombinant AAV, which subsequently was used to infect Cas9-expressing N2A cell line and successfully introduced its 2-piece cargo into the targets achieving desired AA changes (Fig. 3C). Robust gene editing efficiency was observed, with ∼5% screened clones (2 out of 46 by Sanger sequencing) showing bi-allelic knock-in (Fig. 3D), 30% (14 out of 46) with hemizygous knock-in, and the remaining 65% with indels (data not shown).

**Figure 3.**
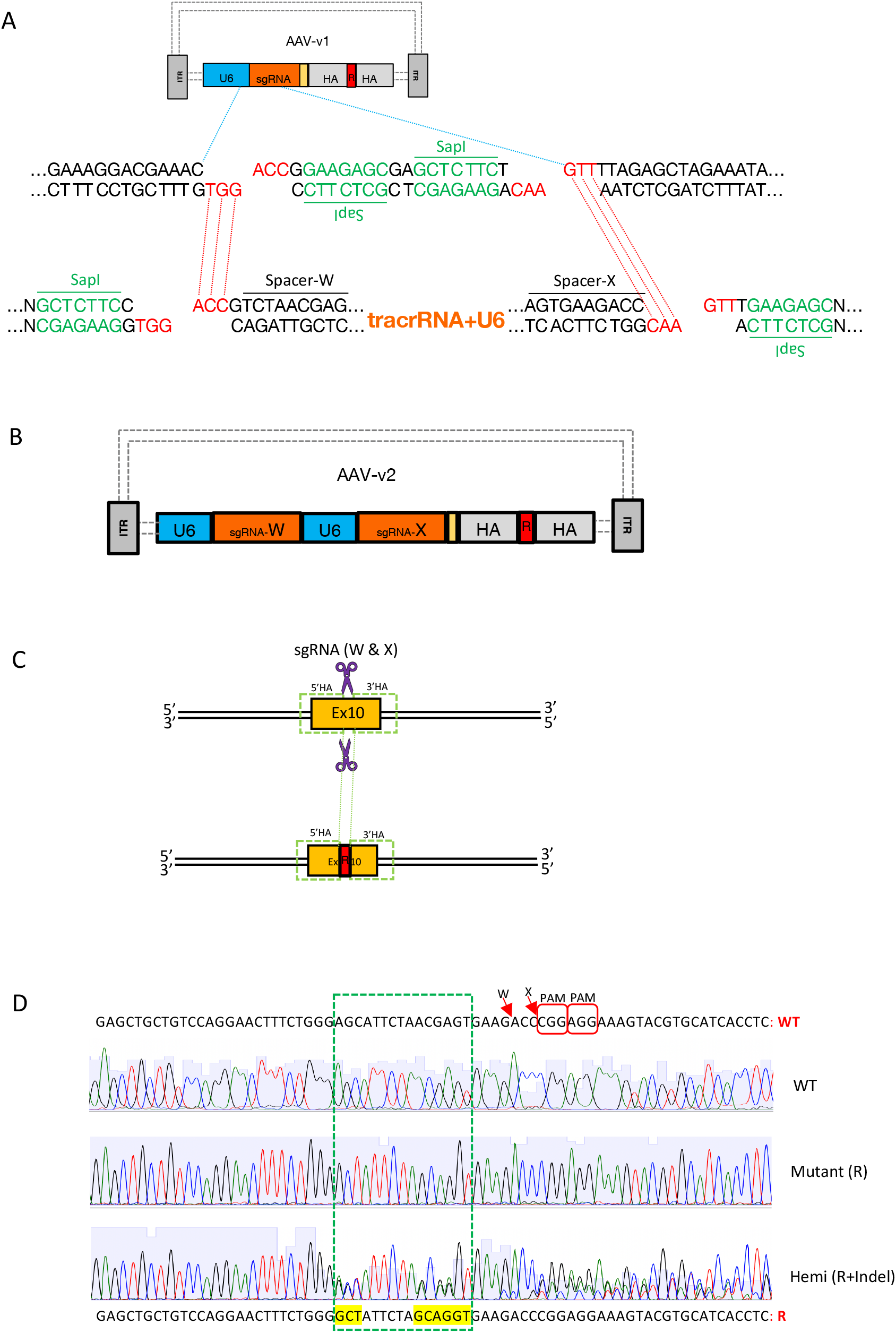
Cloning duo-guides into AAV-v1 vector by exploiting built-in cloning site (originally designed for one sgRNA). Gene editing (AA changes at PSEN1 gene) in N2A cell line. **(A)** (top) Schematic AAV-v1 construct: blue dotted line zooming out partial sequences of U6 and sgRNA cloning site. (middle) Type IIS RE (SapI) digest creates two unique 5’-overhangs (GGT vs. GTT). (bottom) Showing gene synthesized plasmid carrying duo-sgRNA cassette sequence (Spacer-W+tracrRNA+U6+spacer-X) with complementary overhangs, flanked by uniquely positioned SapI sites, which upon SapI digestion renders two complementary 5’-overhangs (ACC vs. AAC; red dotted line) for cloning. **(B)** Schematic of the final construct AAV-v2. **(C)** (top) CRISPR guides (W and X) target sites (purple scissors) on PSEN1 exon 10 region. (bottom) AA replacement donor (R) integrated at the genomic target after rAAV-v2 transduction. **(D)** PSEN1 exon 10 sequence shown (WT vs. R: 3/5 AA changes - three amino acid codons yellow highlighted). Red arrow: SpCas9 guides (W and X) cutting sites; chromatograms showing WT, mutant (R) and hemizygous (R+indel); green dotted rectangle encompassing the 3/5 AA changes within a 15-nt range.

### CRISPR-*CLIP* (CRISPR-*<u>C</u>*lipped *<u>L</u>*ong ssDNA via *I*ncising *<u>P</u>*lasmid): for procuration of lssDNA donor

Various approaches have been proposed to construct lssDNA templates since the introduction of *Easi*-CRISPR, including *iv*TRT (*in vitro* transcription and reverse transcription) from the original *Easi*-CRISPR protocol (19); dsDNA plasmid-retrieval-based method using RE (BioDynamics Laboratory kit); PCR-based method which uses a phosphorylated primer to label the undesired DNA strand for degradation (Takara Bio kit); chemical synthesis by commercial vendors (e.g., Megamer by IDT). These methods enable the procuration of lssDNA with sequence fidelity (except *iv*TRT) and length extension up to ∼2 kbases. Certain constraints, however, could arise from the use of RE (efficacy, availability, or unintended RE cut on the donor sequence) as well as from technical difficulties in PCR-based or chemical synthesis (e.g., complex sequences with unusual repeats or specific nucleotides composed in too high/low percentage) to procure complex sequences, thus failing the lssDNA construct. Alternatively, we devised CRISPR-*CLIP* here to avoid such pitfalls.

To generate the lssDNA donor for creating conditional knock-out (CKO) mouse model of GENE-Y* via *Easi*-CRISPR, we first obtained the gene synthesized dsDNA template which consists of a floxed cassette of exon2, flanked by HA to the genomic target. The template is 2.2 kb in length and anchored in a default plasmid (pUC57) (Fig. 4A). As the HA regions of the donor template encompass sequences with various types of unusual repeats which failed the lssDNA procuration via PCR-based method (data not shown), we adopted CRISPR-*CLIP* instead. We used Cpf1 and Cas9n (D10A mutant nickase version of Cas9) (1, 7, 20) with corresponding guides to induce a dsDNA cleavage and a nick, respectively, on the plasmid at two junction sites flanking the lssDNA cassette; specifically Cas9n was exerted on the strand of interest (lssDNA) (Fig. 4B; guides CLIP-B and CLIP-A; Table S1). The resulting three stand-alone single-stranded DNA (ssDNA) units were of unique sizes hence were able to be separated on agarose gel electrophoresis upon denaturing gel-loading buffer (DGLB) treatment (Fig. 4C). The 2.2-kbase target strand of our interest (i.e. lssDNA donor) was thus identified and we clipped it out through gel extraction procedure.

**Figure 4.**
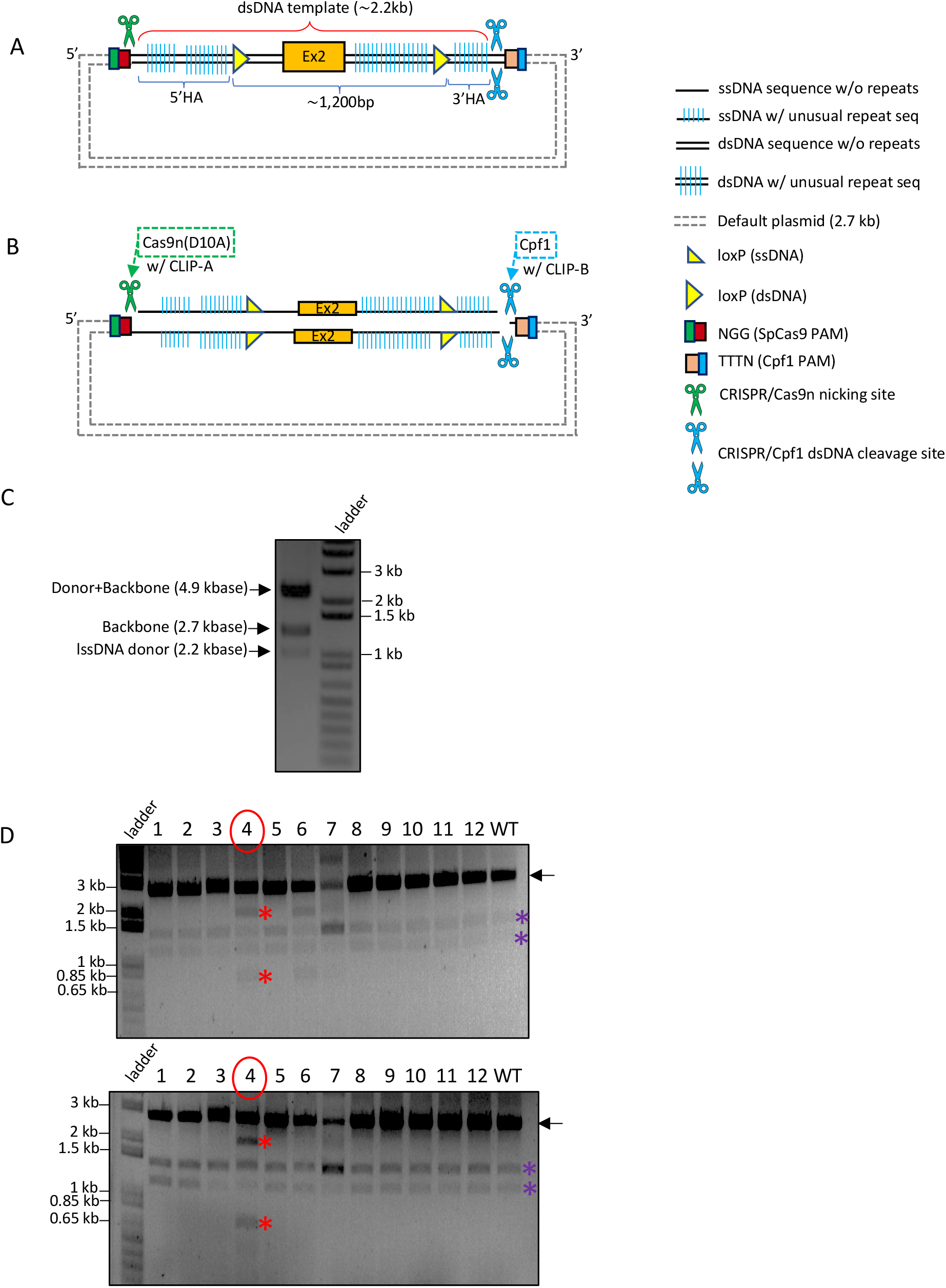
CRISPR-*CLIP*: Procuration of lssDNA from dsDNA template & Genotyping results of the CKO mice generated with the acquired lssDNA via *Easi-CRISPR*. **(A)** dsDNA template anchored in the default plasmid; the sense ssDNA (top strand) is the donor (lssDNA) of choice. **(B)** Cpf1 (with guide CLIP-B) was used to create a dsDNA incision on the plasmid at one end of the lssDNA cassette, while Cas9n (with guide CLIP-A) to create a ssDNA incision at the other end, specifically on the strand of interest (top strand in this case). **(C)** Upon DGLB treatment, the plasmid incised by Cpf1 and Cas9n was resolved into three stand-alone distinct-sized units (0.9% agarose gel electrophoresis): ∼4.9 kbases (donor+backbone) vs. ∼2.7 kbases (backbone) vs. ∼2.2 kbases (lssDNA donor). **(D)** Mice genotyping screened by RE HindIII (top) and EcoRV (bottom): a pair of external screening primers amplified 2.8 kb DNA fragment (black arrow); mice with the lssDNA donor integration should carry a floxed cassette with HindII/EcoRV site adjacent to LoxP. Upon RE digestion, mouse #4 showing the insertion of both LoxPs (∼0.8 kb vs. ∼2 kb; red asterisk), further confirmed by Sanger sequencing; mouse #6 showing only HindIII digest, indicating one LoxP insertion; mouse #7 was found to carry heterozygous 1.2 kb deletion between the two guides (used for creating CKO model), verified by Sanger sequencing, thus showing a band at ∼1.6 kb. Due to incomplete RE digest and usage of EtBr pre-stained gel, the smaller bands appeared in lighter intensity; purple asterisk: non-specific PCR band.

The lssDNA acquired was subjected to validations for its intactness, single strand feature, and sequence integrity. To verify length intactness, the 2.2-kbase lssDNA was resolved on gel electrophoresis in parallel to its counterpart of source dsDNA (2.2 kb) for size comparison. For obtaining the latter as control reference, we used WT Cas9 and Cpf1 to digest the lssDNA-carrying plasmid (comprising 2.2 kb dsDNA version of template, plus 2.7 kb pUC57 backbone, leading to 4.9 kb in total length), which yielded two DNA fragments migrating at respective sizes on the agarose gel (Fig. 5A, lane b); whereas the 2.2 kbase-lssDNA donor migrated in the vicinity of 1.1 kb, indicating intact length (Fig. 5A, lane d). The single-stranded feature of the donor was verified by using dsDNA-specific RE BamHI to digest the lssDNA that carries a BamHI cut site. Upon BamHI digestion (in addition to WT Cas9 and Cpf1), the 4.9 kb lssDNA-carrying plasmid, served as a positive control, resulted in three digested fragments (Fig. 5B, lane c), while the acquired lssDNA did not yield digestion product (Fig. 5B, lane e). Lastly, the sequence integrity was validated by Sanger sequencing using complementary primers (Fig. 5C). As a negative control, non-complementary primers were used to sequence the lssDNA and failed to pick up correct reading, further reflecting the lssDNA purity (not contaminated with dsDNA). The lssDNA yield of ∼50% was obtained (e.g., digesting 100 µg of 2.2 kb dsDNA in 2.7 kb pUC57 backbone would result in >10 µg of lssDNA). The lssDNA thus procured was supplied as the donor into mouse zygotes via pronuclear microinjection and successfully generated CKO mouse model at 8% (1 out of 12 founder mice showed both LoxP inserted) targeting efficiency (Fig. 4D).

**Figure 5.**
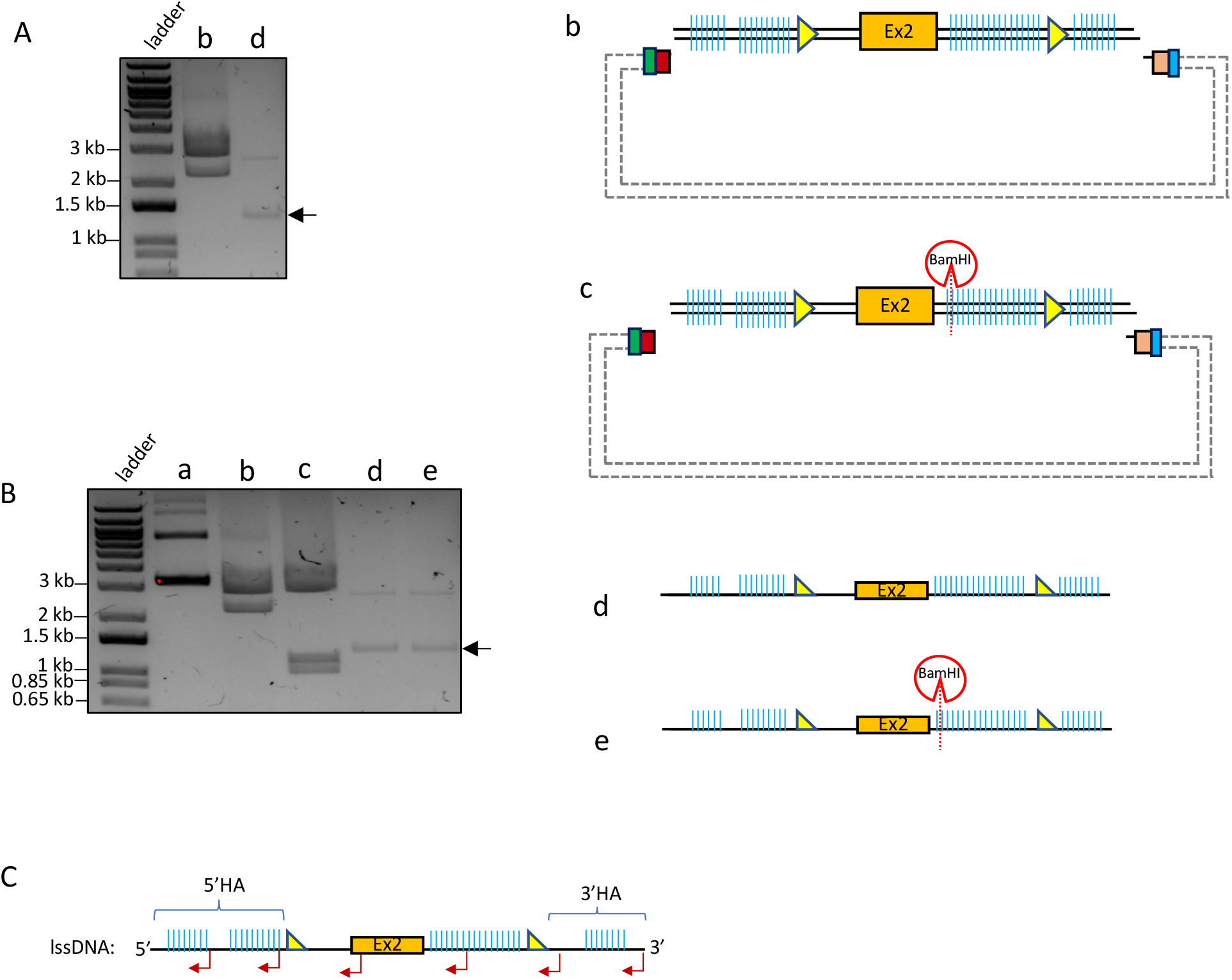
Validation of lssDNA acquired by CRISPR-*CLIP*. **(A)** Cpf1 & WT Cas9 digested the entire plasmid (∼4.9 kb, as shown in Fig. 4A) into ∼2.7 kb and ∼2.2 kb DNA fragments (lane b); acquired lssDNA (∼2.2 kbase) resolved around 1.2-1.3 kb in size^$^ (lane d, arrow). **(B)** Control: the uncut plasmid (lane a); control: as described in (A) (lane b); Cpf1 and WT Cas9, along with additional BamHI digested the plasmid into three fragments: BamHI cleaved the ∼2.2 kb dsDNA template into ∼1.2 kb and ∼1 kb, while ∼2.7 kb default plasmid remained intact (lane c); BamHI digestion did not cleave the lssDNA despite bearing a BamHI cut site (lane e, arrow) vs. lssDNA without BamHI digestion (lane d); all resolve in 0.9% agarose gel electrophoresis. **(C)** Multiple reverse primers (red bent arrow) were used for Sanger sequencing validation of the acquired lssDNA (sense DNA, in this case). ^$^The lssDNA may not resolve at the exact predicted size (1.1 kb in this case), depending on certain factors, such as buffer condition and potential secondary structure formation due to sequence composition, in which case additional DGLB treatment on acquired lssDNA could aid in size separation with more precise outcomes. The integrity of the procured lssDNA is considered of good quality as long as the majority of the lssDNA resolved close to the predicted size.

## Discussion

The two CRISPR-based methods shown in the above three cases offer effective strategies to overcome technical challenges frequently encountered in constructing donor templates for genome editing. Preferring to avert amplification issues with PCR, the CRISPR-*CLONInG*, can be adjusted to utilize CRISPR/Cas9 as the excision tool to also acquire every vector component, rather than only the backbone. In that case, the required Gibson overhangs for facilitating seamless cloning can be restored by additionally supplying a short gBlock carrying complementary sequences to backbone-insert or insert-insert junction for Gibson Assembly reaction.

The CRISPR-*CLIP* demonstrates a PCR-free-and-RE-free strategy that imposes no restrictions on sequence composition for procuring lssDNA donor templates. We have now routinely applied this method to generate donors that have been used for creating novel gene knock-in, humanization, CKO, and more recently for conditional knock-in (CKI) models in mice (Fig. S1A). The latter case carried a Lox66/71 floxed cassette, with PSEN1 WT (exon 10) and the inversely positioned mutant (exon 10 with 15 bp AA replacement sequence) fused within, leading to a template of 3.5 kbases in length (Fig. S1B, S1C, SI Sequence 4). These CRISPR-*CLIP*-mediated donors collectively exemplify an effective approach to procure lssDNA templates encoding fairly large and yet diverse genetic modifications for efficient generation of mouse models via *Easi*-CRISPR. That said, finding the prerequisite PAM for CRISPR-*CLIP* target sites for incisions on the dsDNA template plasmid may not always be possible, especially in Cpf1’s case. To cope with such issue, two add-on sequences of Cpf1-Cas9 (duo-PAM-A and duo-PAM-B) can be placed on the plasmid at the exact junction sites flanking the lssDNA cassette (duo-PAM-A at upstream end; duo-PAM-B at downstream end) (Fig. 6A), which also streamlines the donor design process. Specifically, duo-PAM-A carries both PAMs with a 19 nts assigned Cpf1 spacer sequence (21), where Cpf1 and Cas9’s PAMs are positioned at the 5’ end of the top and bottom strands, respectively, of the add-on DNA-tag; while duo-PAM-B carries both PAM sequences in inverse orientation of the duo-PAM-A’s (Fig. 6B; Table S1). Moreover, both duo-PAM tags are assigned with distinct spacer sequences such that the DNA strand polarity of choice can be acquired by simply choosing suitable Cas type to make exclusive type of incision on the plasmid at the particular end of the lssDNA cassette junction: using Cas9n to incise duo-PAM-A whereas Cpf1 to incise duo-PAM-B for acquiring the top strand (Fig. 6B, 6C) vs. using Cpf1 to incise duo-PAM-A whereas Cas9n to incise duo-PAM-B for acquiring the bottom strand (Fig. 6B, 6D).

**Figure 6.**
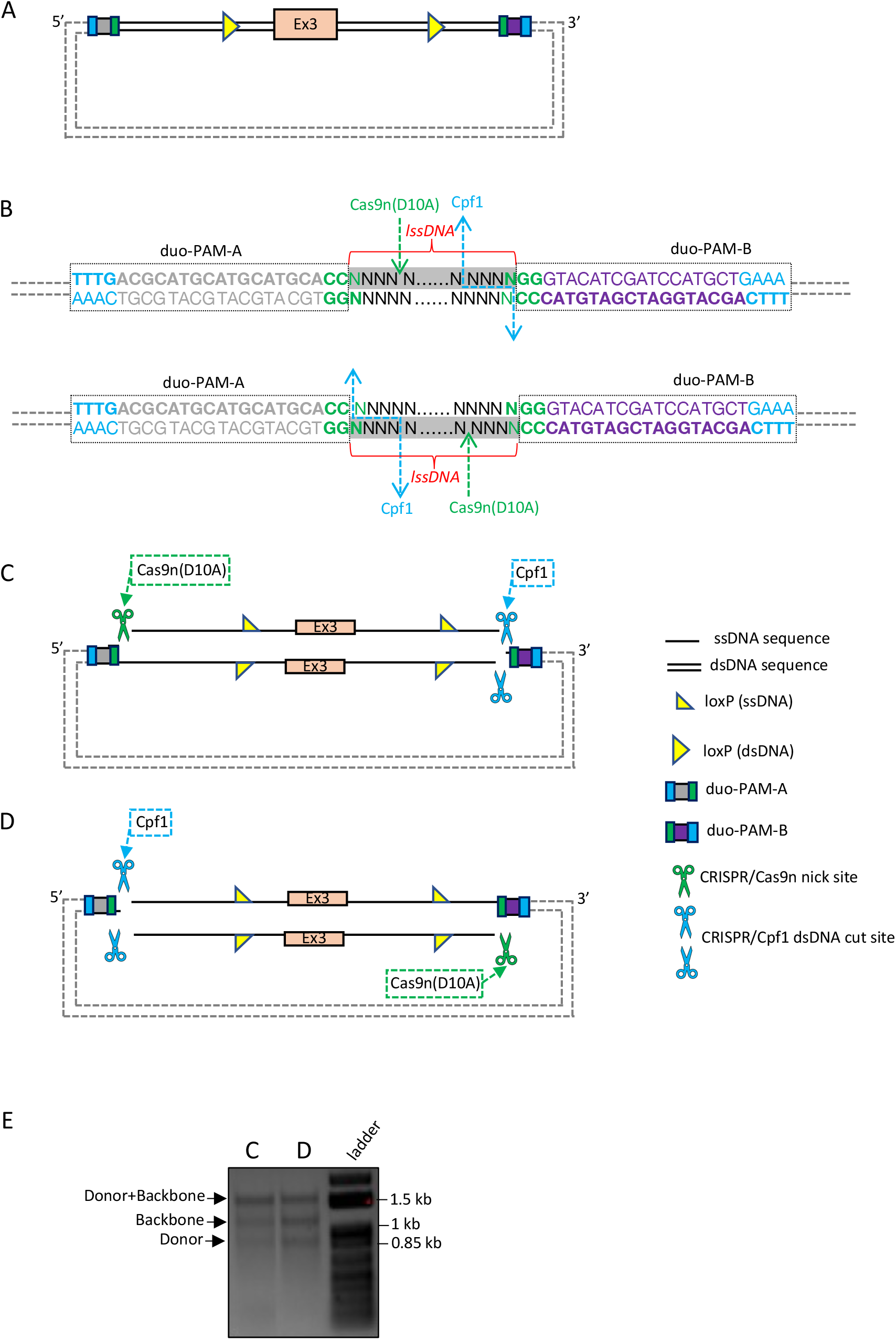
CRISPR-*CLIP* with add-on feature of universal DNA-tag sequences of Cpf1-Cas9 duo-PAM. **(A)** The dsDNA template carrying lssDNA^#^ cassette is flanked with duo PAM-A (upstream end) and duo-PAM-B (downstream end), anchored in the default plasmid. **(B)** Each duo-PAM tag contains 23-bp sequence with PAMs for Cpf1 and Cas9 placed at respective 5’ ends of each DNA strand, plus a constant Cpf1 spacer sequence. Duo-PAM-A and -B are assigned with two distinct Cpf1 spacer sequences to enable suitable Cas types to make exclusive incisions on the plasmid at both ends of lssDNA junction sites. Of note, Cas9 spacer sequence is variable and subject to lssDNA donor; Cpf1 and Cas9 incisions on the duo-PAM tags will each remove 4 nts (end sequences of HA) from the lssDNA donor, which merely results in trivial variation in HA length. **(C)** To procure the sense ssDNA (top strand) as the lssDNA donor, choose Cas9n to cut at duo PAM-A end while choosing Cpf1 to cut at duo PAM-B end. **(D)** To procure the antisense ssDNA (bottom strand) as the lssDNA donor, choose Cpf1 to cut at the duo PAM-A, while using Cas9n to cut at the duo-PAM-B. **(E)** Upon DGLB treatment, C and D respectively yielded 3 stand-alone ssDNA units of distinct sizes resolved in 0.9% agarose gel electrophoresis. ^#^Belongs to a locus different from the case shown in Figures 4 and 5.

Further, the implement of duo-PAM is reasoned to incorporate the provision for top vs. bottom strand choice of the plasmid as a preferential lssDNA donor. A kinetic study reported long residency time of Cas9 on DNA double-stranded break target site with asymmetric dissociation timeline from four broken strands, wherein 3’ end of the cleaved DNA strand that is not complementary to the sgRNA (nontarget strand) is released first while the other three strands still tethered to Cas9-sgRNA complex (22). Such scenario implicates accessibility lag among the cleaved DNA strands for initiating strand complementation in the DNA repair process. We hypothesize that supplying a lssDNA donor that carries complementary sequence to the nontarget strand exploiting its PAM-distal (3’) end with immediate accessibility may potentially gain leverage in the recombination event, thereby enhancing the donor insertion rates; in this same context, extending the HA that is homologous to the PAM-proximal end longer than the PAM-distal end complementary HA, aimed to offset possible ssDNA exonuclease degradation of the former HA. While this hypothesis warrants further investigation, the add-on duo-PAM feature provides feasibility for choosing the optimal strand of lssDNA donor when suitable, regardless of availability of pre-existing PAM(s) in the plasmid for CRISPR incision(s). Overall, the CRISPR-*CLIP* with duo-PAM add-on further simplifies the lssDNA generation process and broaden its applicability to suit any genomic sequences of interest, and together with CRISPR-*CLONInG*, both methods illustrate strategies for efficient construction of highly customizable donor templates for facilitating precision genome engineering.

## Methods and Materials

### CRISPR digestion of vector backbone for CRISPR-*CLONInG*

To form ctRNP complex of Cas9 (IDT, cat#1081058):crRNA (IDT):tracrRNA (IDT, cat# 1072532) in 1:2:2 ratio, 48.8 pmole of ctRNA (crRNA and tracrRNA) was added in a RNAse-free tube, heated at 100 ℃ for 2 min, and cooled at RT for 10 min. Cas9 protein (2 µg) was added and incubated at 37 ℃ for 10 min to form the ctRNP complex. About 2-3 µg of source DNA plasmid (<10 kb) was added with NEBuffer 3.1 (NEB, cat# B7203S), DEPC ddH_2_O in a volume of 30 *μ*l and incubated at 37 ℃ for >2 hour (h). Cas9 was inactivated at 70 ℃ for 15 min, ctRNA was degraded with 10 µg of RNase A (Qiagen, cat#19101) at 37 ℃ for 30 min and resolved on a 0.9% agarose gel electrophoresis, followed by gel extraction to acquire vector of interest.

### PCR amplification of vector insert for CRISPR-*CLONInG*

All the primers (Table 2) were ordered from IDT and Eurofins Genomics. Primer pair Neo-F & Neo-R was used to amplify FRT-Neo-FRT from pL451 plasmid (23); tdTom-F & tdTom-R was used to amplify tdTomato gene from an existing plasmid. Primer pair AAV-F & AAV-R was used on custom gene synthesized dsDNA anchored in a default vector that carries 15 bp knock-in sequence flanked with two HA (∼400 bp each). Two different PCR systems were adopted: Herculase II Fusion DNA polymerase (Agilent, part# 600679), and Accuprime Pfx DNA polymerase PCR system (Thermofisher Scientific, cat# 12344-024). The PCR condition was 95 ℃ for 3 min; 30 cycles of 95 ℃ for 30 sec, 60 ℃ for 30 sec, 72 ℃ for 1 min/kb; 72 ℃ for 5 min. PCR amplified DNA fragments were subjected to 0.9% agarose gel electrophoresis and DNA of expected size were gel purified.

### Gibson (HiFi) Assembly for CRISPR-*CLONInG* and Ligation reaction

The CRISPR digested vector backbones (∼50 ng for each FLEx and AAV) and PCR-amplified inserts carrying Gibson overhangs (FRT-Neo-FRT and tdTomato for FLEx; novel donor template for AAV) were assembled in a ratio of 1:2 with Gibson (HiFi) DNA Assembly Master Mix (NEB, cat# E2621S) following manufacturer’s protocol. The reaction mix was incubated at 50 ℃ for 1 h, and 2 µl of the assembled mix (∼5 ng of vector backbone) was transformed into competent cells (NEB, cat# C2987H), followed by spreading on antibiotic-selective LB agar plates. Mini-prep (Qiagen, cat# 27104) DNA was digested with appropriate RE(s) for diagnostic test followed by Sanger sequencing validation. For cloning of duo-guides for AAV-v2 assembly via standard ligation reaction, T4 DNA ligase (NEB, cat#M0202) was used according to the manufacturer’s protocol.

### CRISPR-*CLIP*

The source plasmid where lssDNA will be retrieved from, as illustrated in Fig. 4A, carries the lssDNA donor cassette (∼2.2 kb) that consists of 5’HA, exon2-floxed sequence, and 3’HA (obtained via gene synthesis from Genewiz). The beginning of 5’HA and towards the end of 3’HA contain PAM sequences for Cas9 and Cpf1, respectively, hence the plasmid was digested at these two sites with Cas9n(D10A) (IDT, cat#1081062) and Cpf1 (IDT, cat#10001272) accordingly. The RNA-guided Cas9 protein used for DNA incision was prepared in 1:2:2 ratio of protein:crRNA:trRNA. Specifically, gRNA for Cas9n (noted as ctRNA_Cas9n(D10A)_) was prepared by mixing 610 pmole of crRNA and 610 pmole of tracrRNA; whereas 625 pmole for Cpf1 gRNA (noted as crRNA_Cpf1_); both gRNA were heated at 100℃ for 2 min, and cooled at RT for 10 min. Cas9n(D10A) and Cpf1, 50 µg each, were added into corresponding gRNA and incubated at 37 ℃ for 10 min to form the protein-gRNA complexes, noted as ctRNP_Cas9n(D10A)_ (<u>ctRN</u>A<u>_Cas9n(D10A_</u>_)_ + Cas9n <u>P</u>rotein) and cRNP_Cpf1_(<u>c</u>r<u>RN</u>A<u>_Cpf1_</u> + Cpf1 <u>P</u>rotein), followed by gently mixing with donor plasmid (100 µg), with NEBuffer 3.1 and DEPC dH_2_O in a volume of 200 µl. The CRISPR digestion reaction was incubated at 37 ℃ for at least 2 h or overnight for better DNA incisions.

Three incisions-bearing plasmid (∼10 µg) was column purified (Invitrogen, cat#K220001) to check for digestion; 1 µg, 2 µg and 3 µg of the eluate, was mixed with 3-fold of Denaturing gel-loading buffer (DGLB) (Diagnocine, cat# DS611), and subjected to 70 ℃ for 5 min, flash cooled on ice for 1min, and resolved in 0.9% agarose gel electrophoresis at constant 100 volts until desired distance is attained. Double digested sample by Cpf1 & WT Cas9 was also included for control reference. Once the lssDNA of interest was separated indicating successful digestion on the agarose gel that requires at least 30 min staining with EtBr (>0.5 *μ*g/ml), the remaining 90 µg of the digested plasmid was DNA precipitated following standard protocol. The 2 µg/lane which gave the best separation was scaled up for extraction using QIAquick Gel Extraction Kit (Qiagen cat# 28704).

### Validation of lssDNA

The dsDNA plasmid (carrying donor cassette) (∼400 ng) was cleaved with ctRNP_Cas9(WT)_ and cRNP_Cpf1_ (∼3 pmole of protein and ∼6 pmole of gRNA) at 37 ℃ for >2 h, followed by inactivation of protein and gRNA degradation following the same method described in the CRISPR-*CLONInG* section. In another replicate reaction, only an extra 20U of BamHI (NEB, cat# R0136S) was added. The CRISPR cleaved +/-BamHI digested samples, along with lssDNA (∼200 ng) +/-BamHI were subjected to 0.9% agarose gel electrophoresis at constant 100 volt. The sense lssDNA (top strand) was sequenced with reverse primers.

### Gene Editing in N2A cell line

N2A cell (Neuro-2a/Cas9-Rosa26-Neo, GeneCopoeia, cat# SL511) was cultured in medium, comprising 44% DMEM (ATCC, cat# 30-2002), 50% Opti-MEM (Life Technologies, cat# 51985-034) with 5% FBS (Gemini, cat# 100-500) and 1% Penicillin-Streptomycin (Millipore Sigma, cat# P4333) in a 37 ℃ humid incubator with 5% CO_2_. The AAV DJ serotype packaging was performed by a vendor (Vigene Biosciences) and obtained a titer of 1.7×10^13^ GC/mL. Overnight rAAV infection in 1:50 ratio was used for optimal editing. The rAAV infected cells were single-cell sorted using the BD FACSAria at our university’s Flow Cytometry Resource Center, and clonally expanded. Genomic DNA was extracted using High Pure PCR Template Preparation Kit (Roche, ref#11 796 828 001), and Herculase II PCR system was used to amplify 1,352bp of exon 10 mutant sequence along with neighboring sequence using TD PCR condition of 95 ℃ for 3 min; 95 ℃ for 15 sec, 61 ℃ for 20 sec (−0.5/cycle), 72 ℃ for 1.5 min, for 10 cycles; 95 ℃ for 15 sec, 56 ℃ for 20 sec, 72 ℃ for 1.5 min, for 25 cycles with primer pair (PSEN1-F & PSEN1-R, Table 2), followed by Sanger sequencing read with PSEN1-seq primer (Table 2).

## Acknowledgements

We thank Jing Gao and Chiayun Han for assistance with mESCs culture and handling; Transgenic and Reproductive Technology Center for zygote microinjection and mouse husbandary; Gali Umschweif Nevo for the gift of Neuro-2a/Cas9-Rosa26-Neo cell line; Daniel Weinberg for providing the original FLEx vector; Flow Cytometry Resource Center for clonal isolation of the N2A cell; Betty Shih of Enlightagen Consulting for help with the manuscript. Funding for this work was provided by The Rockefeller University through the CRISPR and Genome Editing Resource Center (DS, CY, VK, PK) and Fisher Center for Alzheimer’s Research Foundation (VB).

## Author Contributions

Designed and conceived the experiments: DS; performed the experiments: DS VK PK; analyzed the data: DS CY VK VB; contributed reagents/materials/analysis tools: DS CY VB; wrote the paper: DS.

## Supplementary Figure Legend

**Figure S1.**
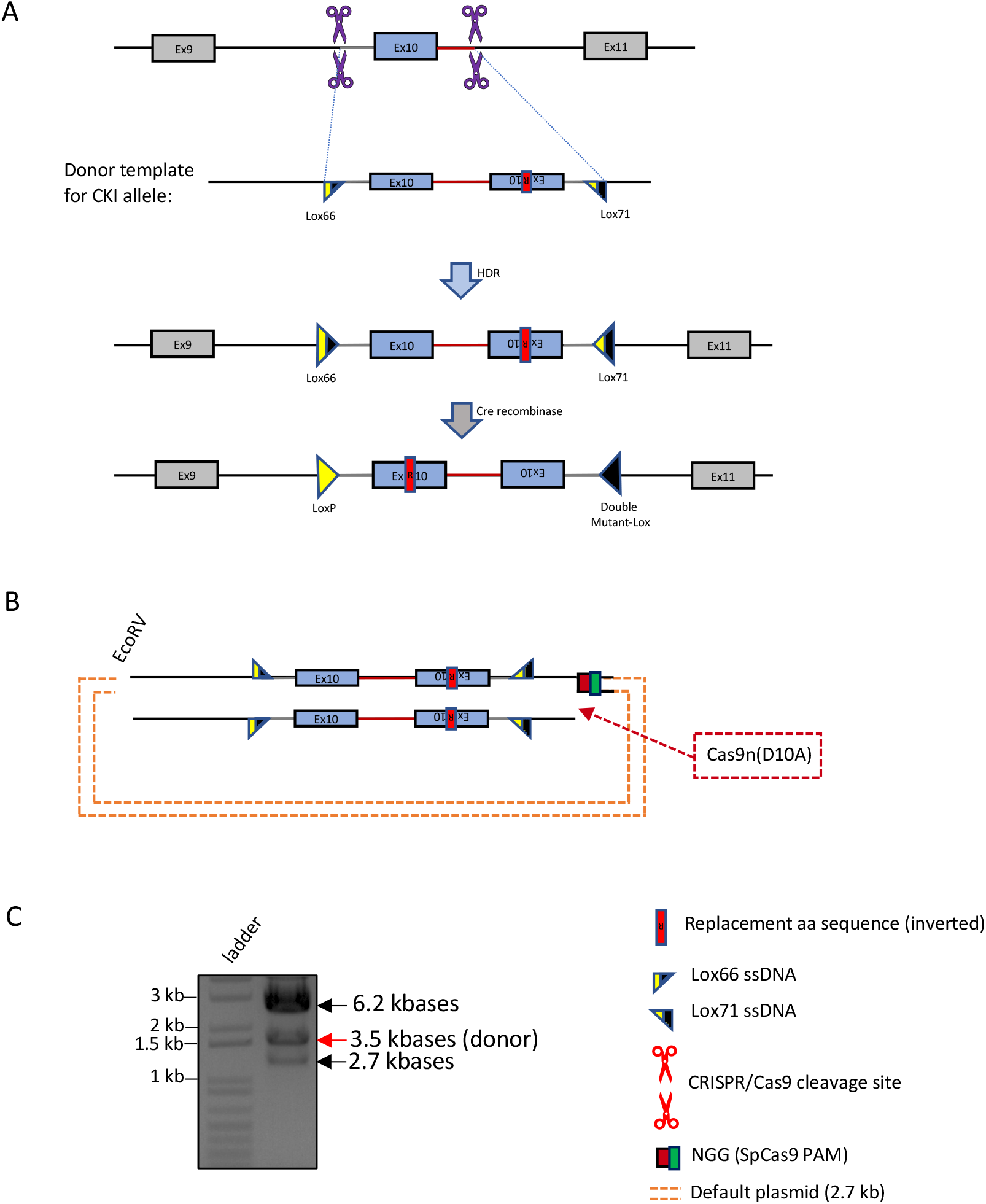
CRISPR-*CLIP*: Using Cas9n and RE (substitute for Cpf1) to generate 3.5-kbase lssDNA. **(A)** Schematic illustration of generating PSEN1 conditional knock-in (CKI) allele. The donor template was designed to carry exon 10 and inverse mutant exon 10, flanked with Lox66 and Lox71 orientated in head-to-head position; once the donor is integrated into the target genome, the expression of Cre-recombinase catalyzes donor inversion to spatio-temporally express the mutant exon 10. **(B)** Plasmid containing assembled dsDNA template anchored in default backbone was incised by Cas9n and EcoRV with ssDNA and dsDNA cleavages, respectively, at respective ends of the dsDNA template. **(C)** The resulting 3 freestanding ssDNA units were treated with DGLB, followed by strand separation on a 0.9% agarose gel (arrows). The strand of interest (3.5-kbase lssDNA donor) was then clipped out via gel extraction and validated (data not shown).

## Supplementary Information (SI)

**Table S1.**
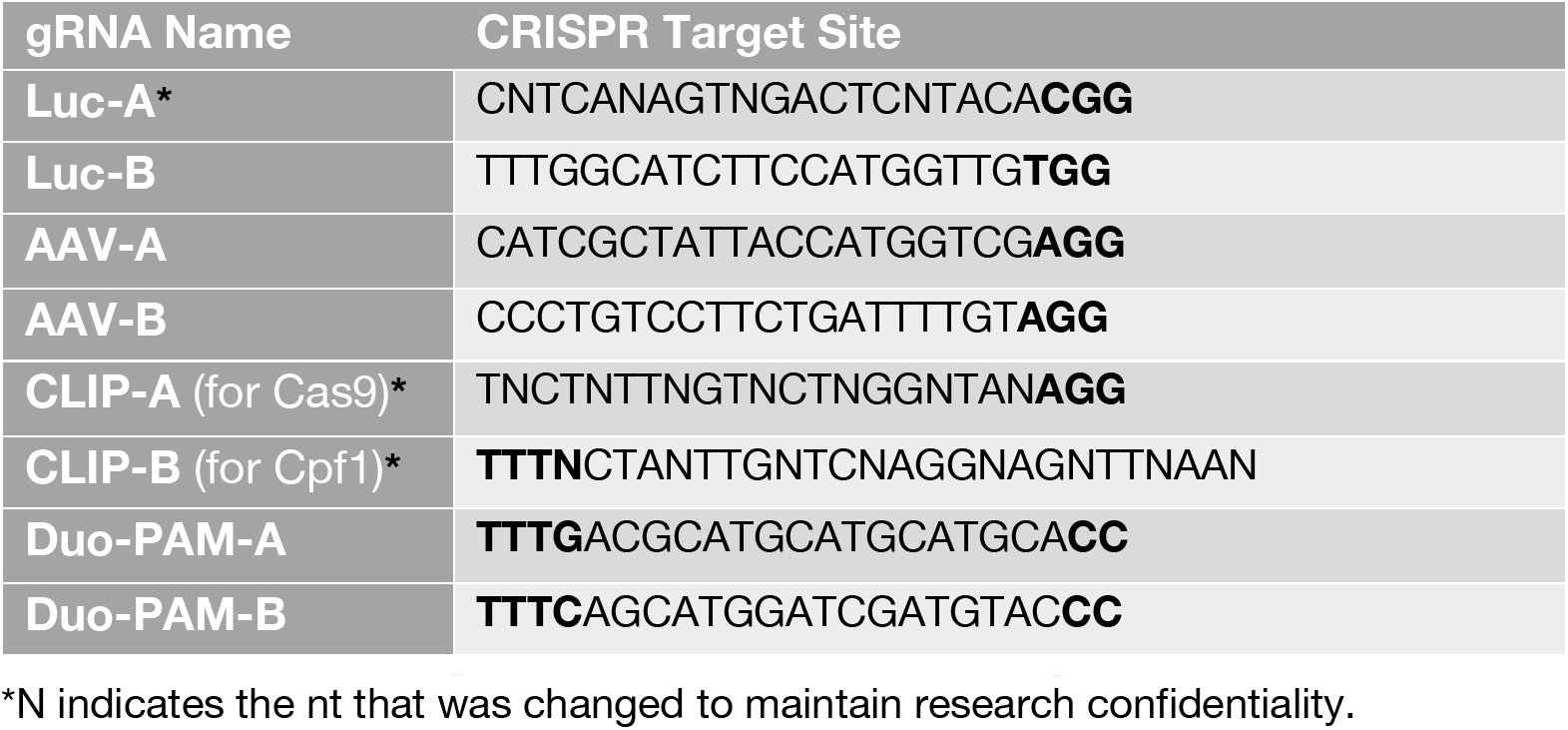
CRISPR guide RNA and their binding sites. PAM bolded.

**Table S2.**
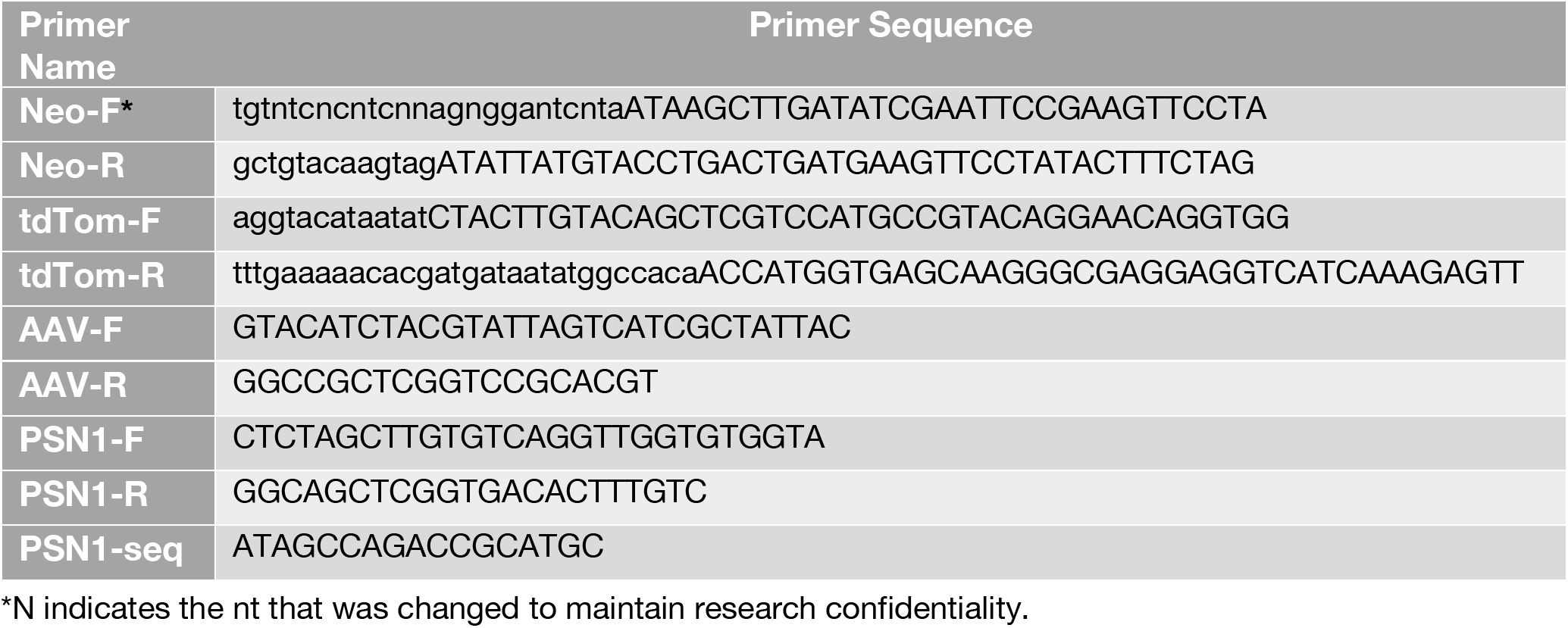
Primers used for PCR amplification of vector inserts from existing plasmids (pL451, tdTomato, AAV from Addgene plasmid# 60229), as well as screening PSEN1 KI in N2A cell line.

**SI Sequence 1.**
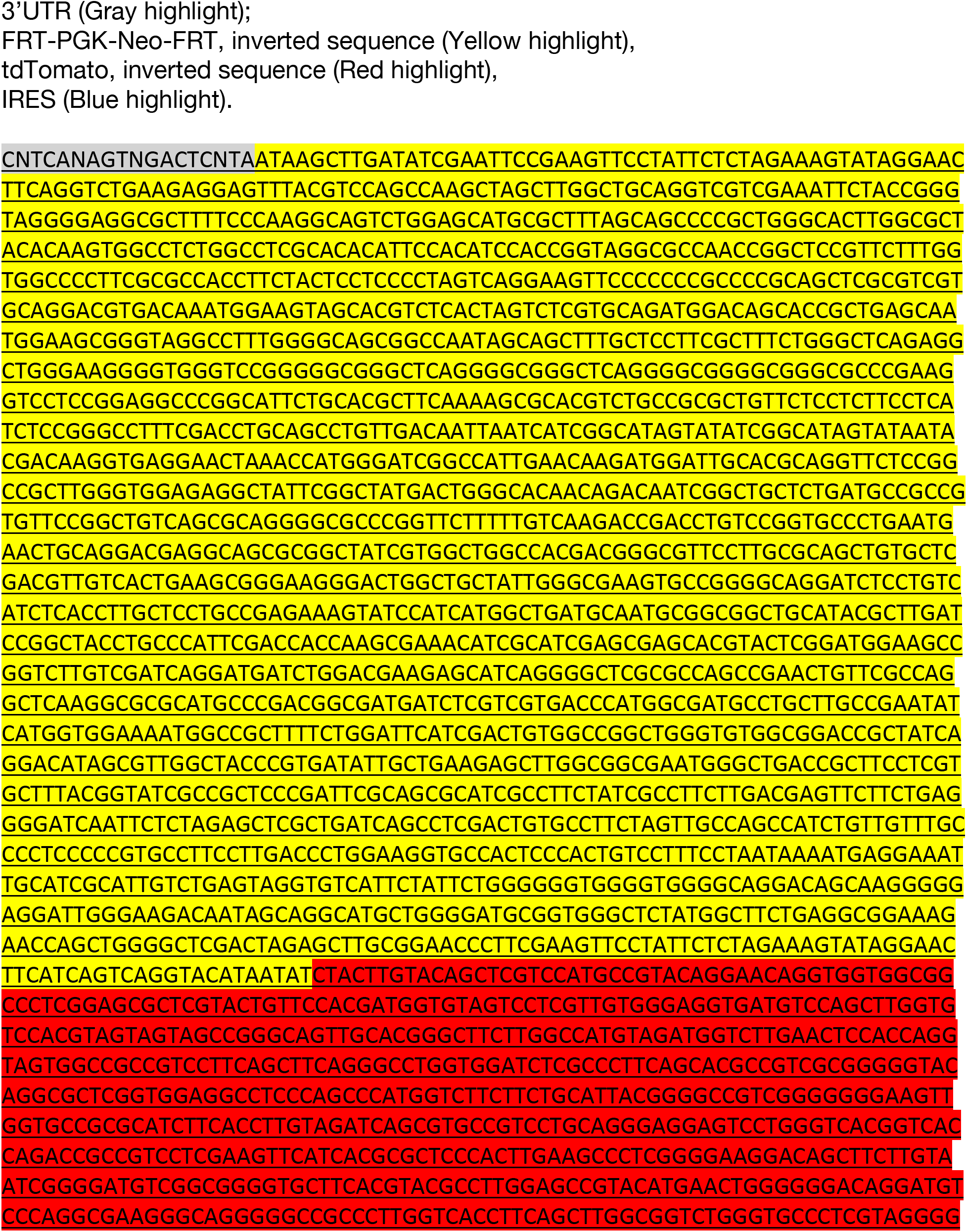
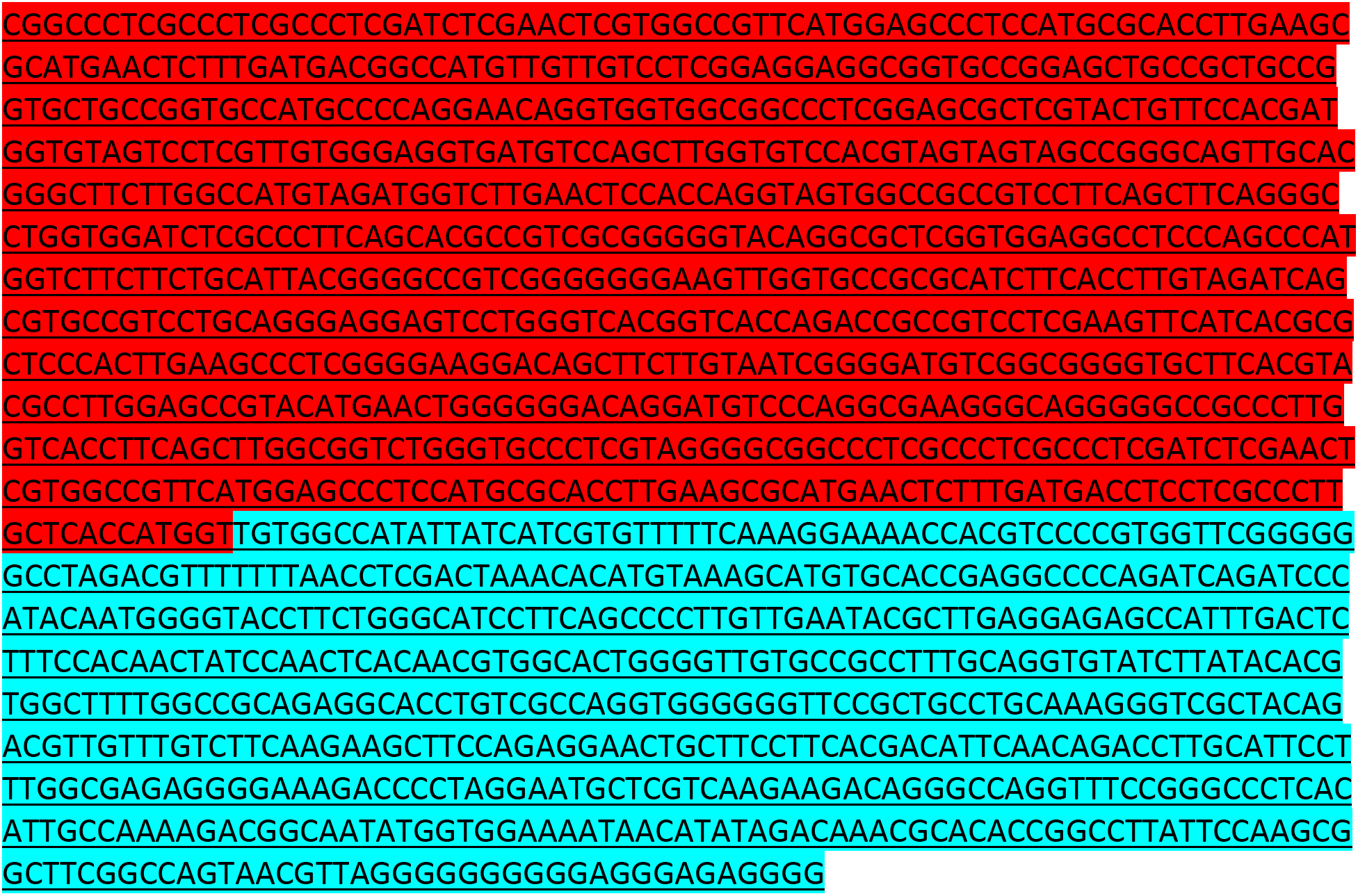
CRISPR-*CLONInG*: Partial sequence of final FLEx vector with partial 3’UTR, FRT-PGK-Neo-FRT, tdTomato and IRES sequences.

**SI Sequence 2.**
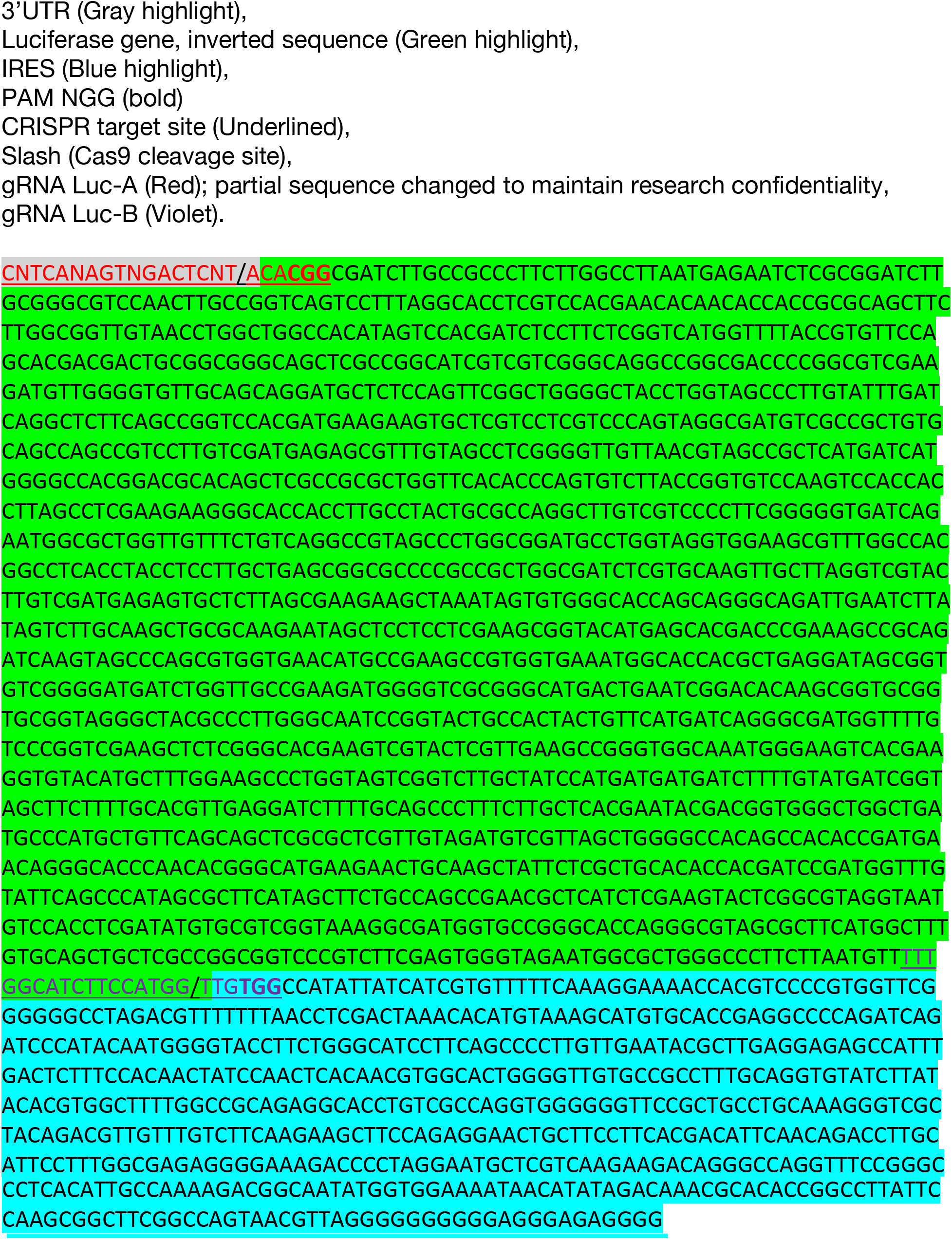
CRISPR-*CLONInG*: Partial FLEx vector sequence showing CRISPR/Cas9 allows precise excision of Luciferase gene from its adjacent 3’UTR and IRES sequences, with only 1 nt deviation.

**SI Sequence 3.**
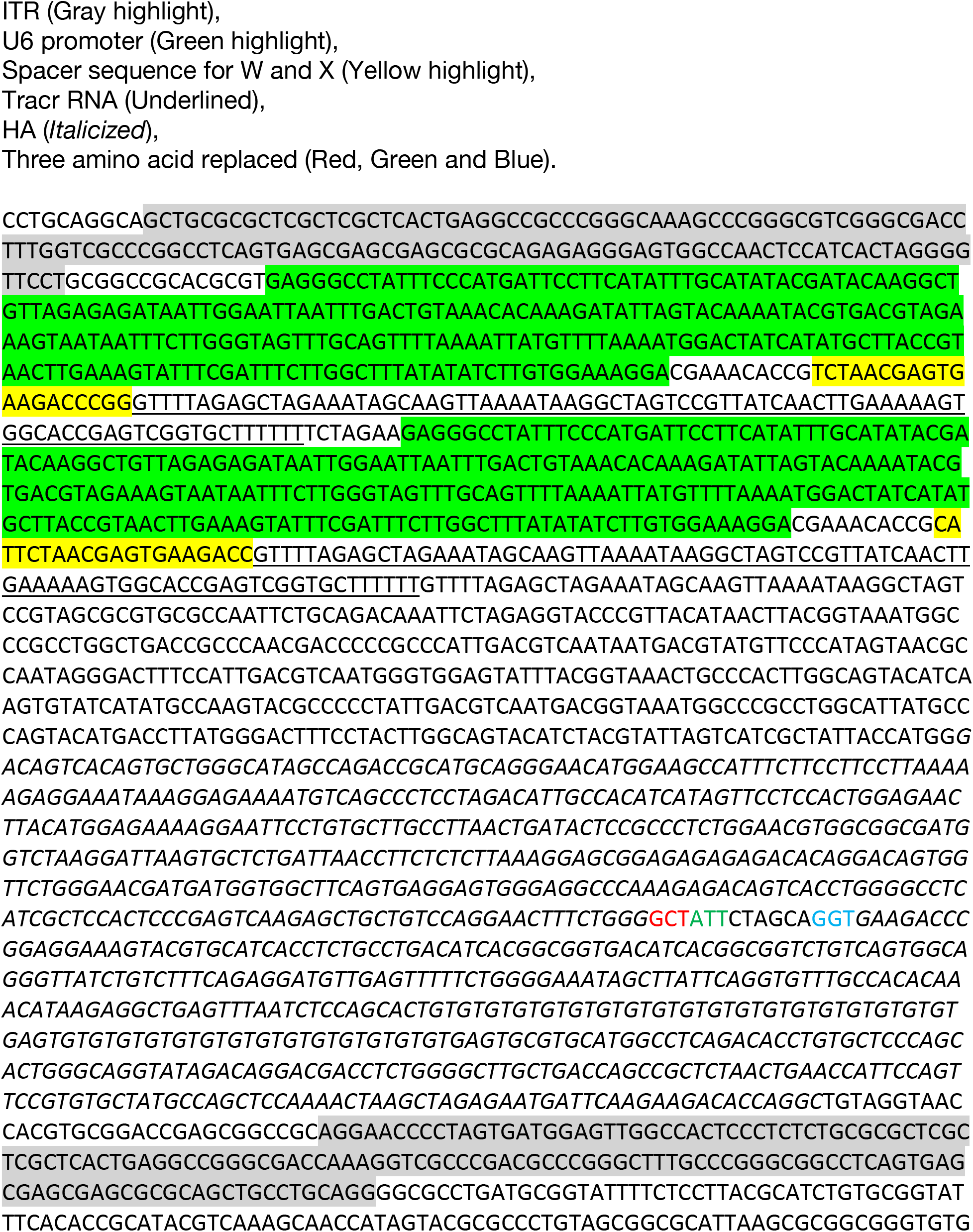
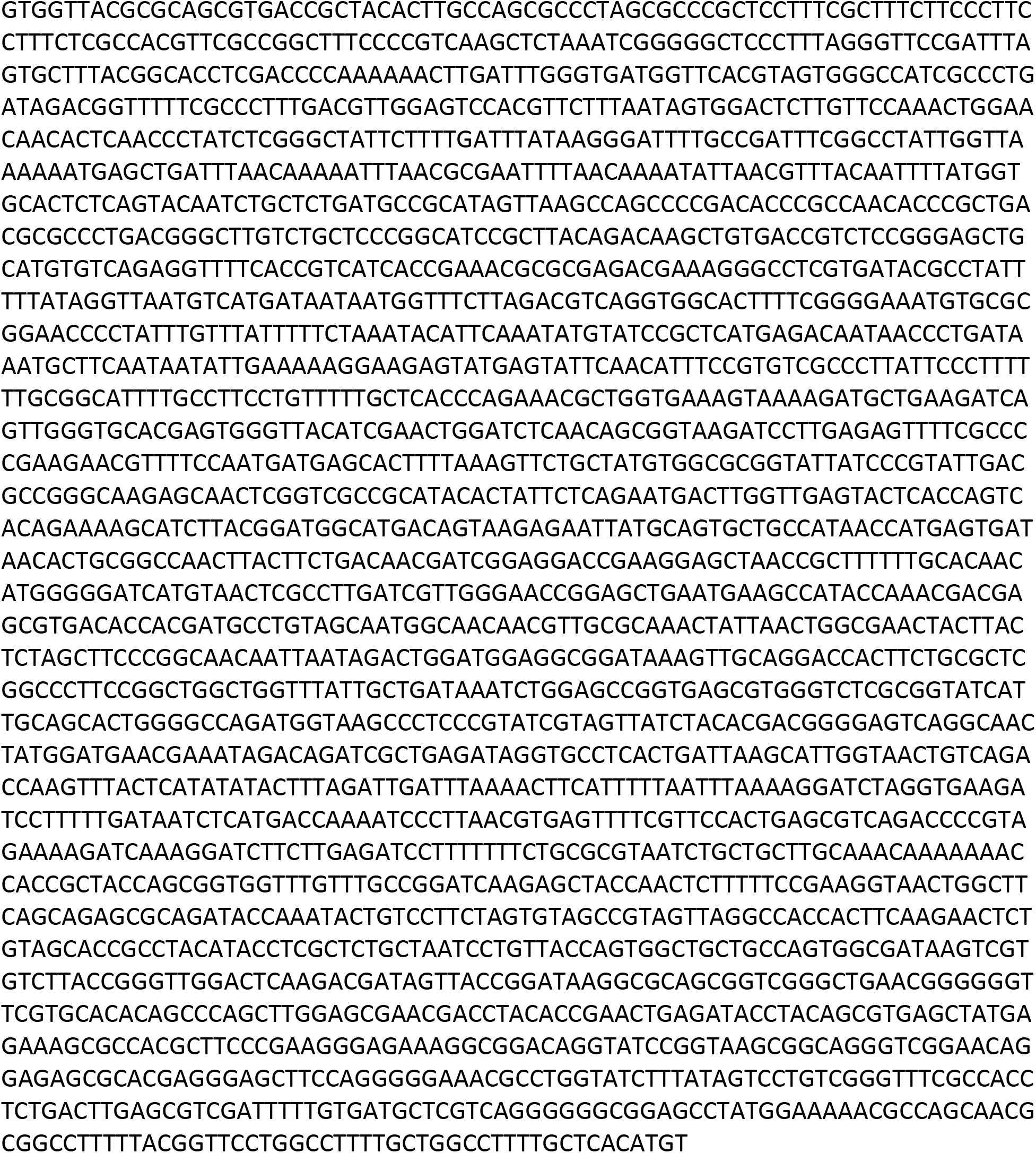
CRISPR-*CLONInG*: Sequence of assembled custom (AAV-v2), comprising sgRNA-W, sgRNA-X, and mutant exon10 of PSEN1.

**SI Sequence 4.**
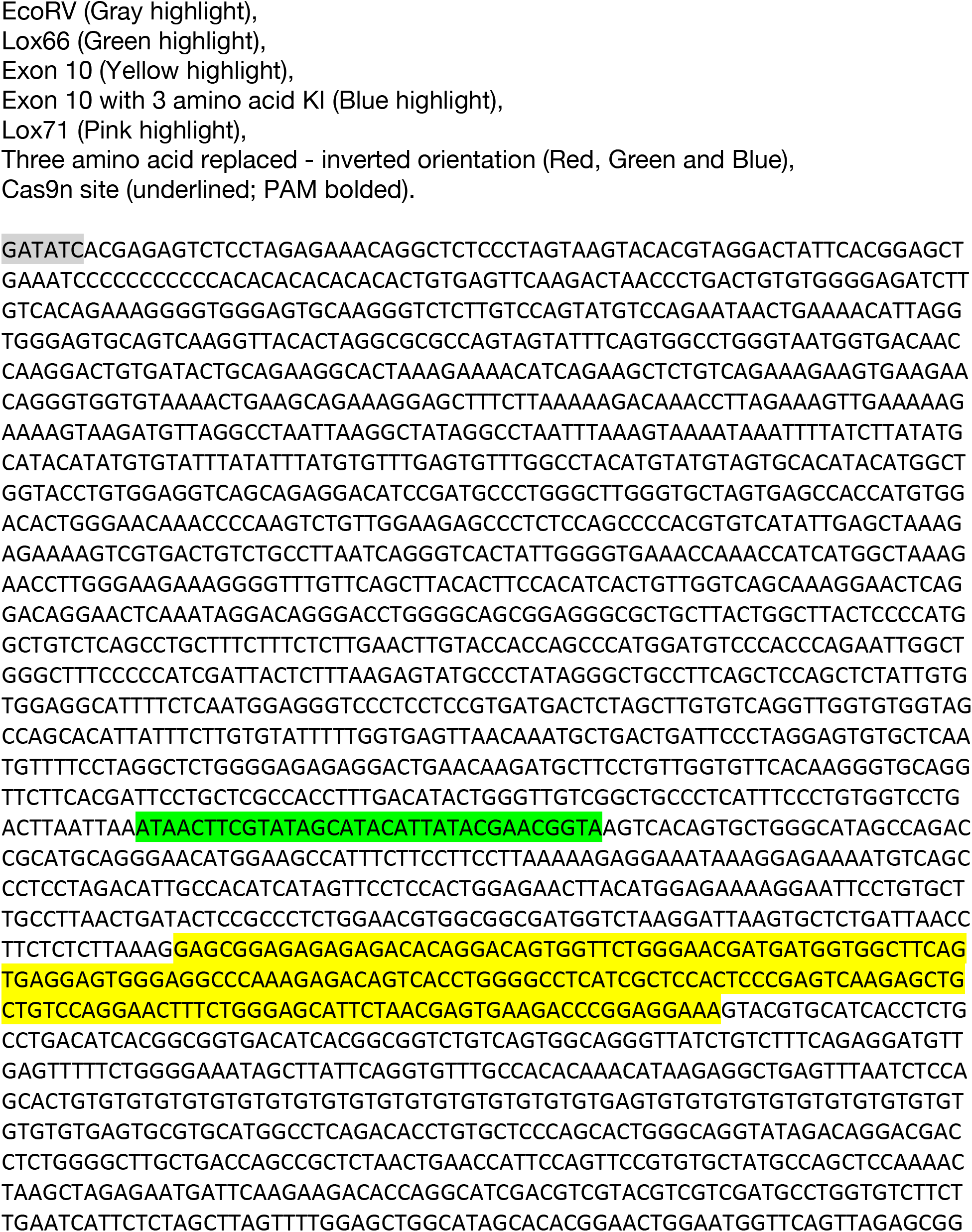
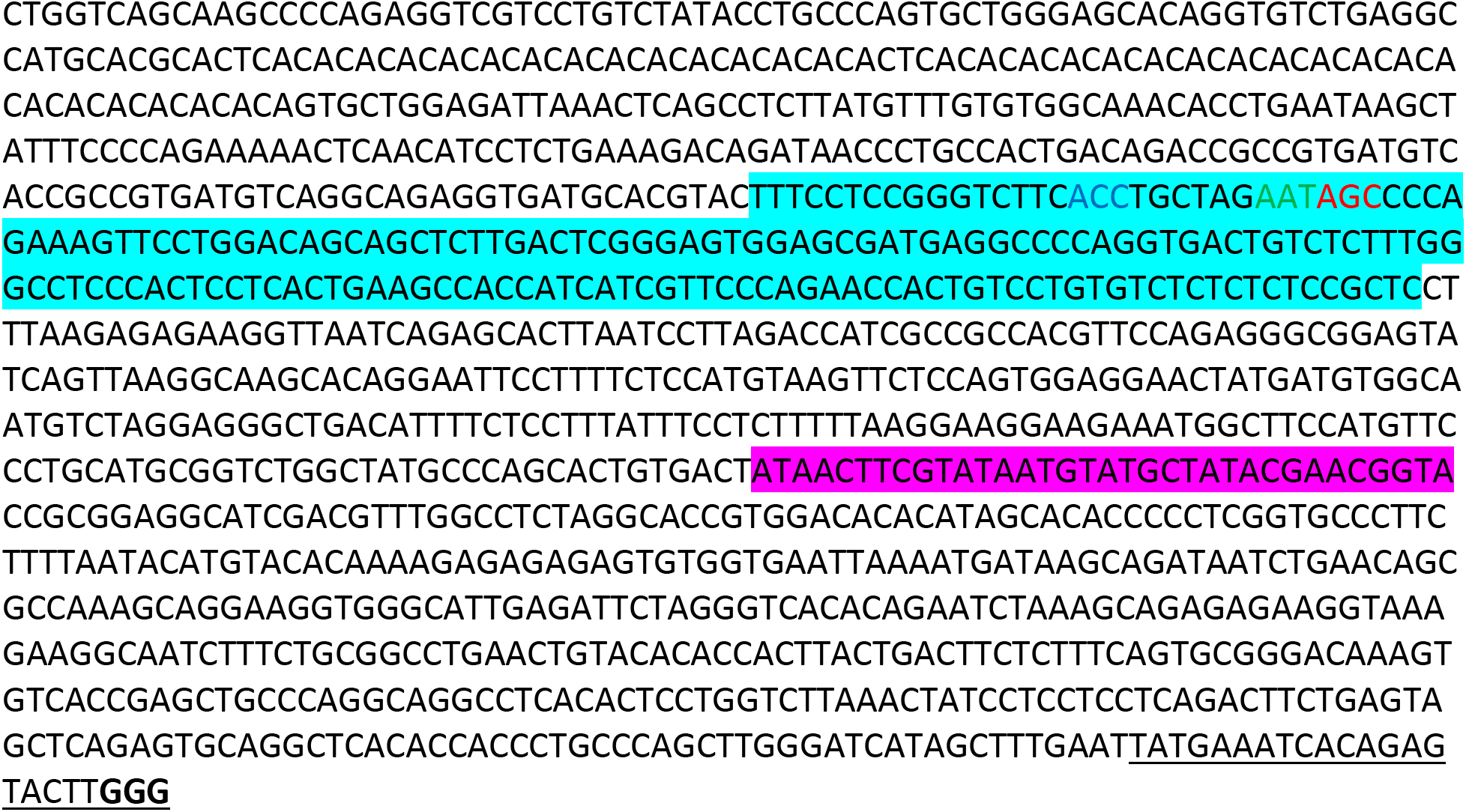
CRISPR-*CLIP*: Sequence of lssDNA donor source template (dsDNA) for PSEN1-CKI allele.

## Footnotes

* To maintain confidentiality of the research, the name of the gene was altered.

## References

1. Jinek M, Chylinski K, Fonfara I, Hauer M, Doudna JA, Charpentier E. A programmable dual-RNA-guided DNA endonuclease in adaptive bacterial immunity. Science. 2012;337(6096):816–21.

2. scherer S. A short guide to the human genome. Cold Spring Harbor Laboratory Press 2008;xiv.

3. Hu JH, Miller SM, Geurts MH, Tang W, Chen L, Sun N, et al. Evolved Cas9 variants with broad PAM compatibility and high DNA specificity. Nature. 2018;556(7699):57–63.

4. Zetsche B, Gootenberg JS, Abudayyeh OO, Slaymaker IM, Makarova KS, Essletzbichler P, et al. Cpf1 is a single RNA-guided endonuclease of a class 2 CRISPR-Cas system. Cell. 2015;163(3):759–71.

5. Sternberg SH, Redding S, Jinek M, Greene EC, Doudna JA. DNA interrogation by the CRISPR RNA-guided endonuclease Cas9. Nature. 2014;507(7490):62–7.

6. Mali P, Yang L, Esvelt KM, Aach J, Guell M, DiCarlo JE, et al. RNA-guided human genome engineering via Cas9. Science. 2013;339(6121):823–6.

7. Cong L, Ran FA, Cox D, Lin S, Barretto R, Habib N, et al. Multiplex genome engineering using CRISPR/Cas systems. Science. 2013;339(6121):819–23.

8. Yang H, Wang H, Shivalila CS, Cheng AW, Shi L, Jaenisch R. One-step generation of mice carrying reporter and conditional alleles by CRISPR/Cas-mediated genome engineering. Cell. 2013;154(6):1370–9.

9. Quadros RM, Miura H, Harms DW, Akatsuka H, Sato T, Aida T, et al. Easi-CRISPR: a robust method for one-step generation of mice carrying conditional and insertion alleles using long ssDNA donors and CRISPR ribonucleoproteins. Genome Biol. 2017;18(1):92.

10. Yoshimi K, Kunihiro Y, Kaneko T, Nagahora H, Voigt B, Mashimo T. ssODN-mediated knock-in with CRISPR-Cas for large genomic regions in zygotes. Nat Commun. 2016;7:10431.

11. Roth TL, Puig-Saus C, Yu R, Shifrut E, Carnevale J, Li PJ, et al. Reprogramming human T cell function and specificity with non-viral genome targeting. Nature. 2018;559(7714):405–9.

12. Gibson DG, Young L, Chuang RY, Venter JC, Hutchison CA, 3rd, Smith HO. Enzymatic assembly of DNA molecules up to several hundred kilobases. Nat Methods. 2009;6(5):343–5.

13. Naso MF, Tomkowicz B, Perry WL, 3rd, Strohl WR. Adeno-Associated Virus (AAV) as a Vector for Gene Therapy. BioDrugs. 2017;31(4):317–34.

14. Vasileva A, Jessberger R. Precise hit: adeno-associated virus in gene targeting. Nat Rev Microbiol. 2005;3(11):837–47.

15. Kohli M, Rago C, Lengauer C, Kinzler KW, Vogelstein B. Facile methods for generating human somatic cell gene knockouts using recombinant adeno-associated viruses. Nucleic Acids Res. 2004;32(1):e3.

16. Miller DG. AAV-mediated gene targeting. Methods Mol Biol. 2011;807:301–15.

17. Platt RJ, Chen S, Zhou Y, Yim MJ, Swiech L, Kempton HR, et al. CRISPR-Cas9 knockin mice for genome editing and cancer modeling. Cell. 2014;159(2):440–55.

18. Swiech L, Heidenreich M, Banerjee A, Habib N, Li Y, Trombetta J, et al. In vivo interrogation of gene function in the mammalian brain using CRISPR-Cas9. Nat Biotechnol. 2015;33(1):102–6.

19. Miura H, Quadros RM, Gurumurthy CB, Ohtsuka M. Easi-CRISPR for creating knock-in and conditional knockout mouse models using long ssDNA donors. Nat Protoc. 2018;13(1):195–215.

20. Gasiunas G, Barrangou R, Horvath P, Siksnys V. Cas9-crRNA ribonucleoprotein complex mediates specific DNA cleavage for adaptive immunity in bacteria. Proc Natl Acad Sci U S A. 2012;109(39):E2579–86.

21. Kleinstiver BP, Tsai SQ, Prew MS, Nguyen NT, Welch MM, Lopez JM, et al. Genome-wide specificities of CRISPR-Cas Cpf1 nucleases in human cells. Nat Biotechnol. 2016;34(8):869–74.

22. Richardson CD, Ray GJ, DeWitt MA, Curie GL, Corn JE. Enhancing homology-directed genome editing by catalytically active and inactive CRISPR-Cas9 using asymmetric donor DNA. Nat Biotechnol. 2016;34(3):339–44.

23. Liu P, Jenkins NA, Copeland NG. A highly efficient recombineering-based method for generating conditional knockout mutations. Genome Res. 2003;13(3):476–84.

